# Inhibition of mitochondrial fission preserves photoreceptors after retinal detachment

**DOI:** 10.1101/197467

**Authors:** Xiangjun She, Xinmin Lu, Tong Li, Junran Sun, Jian Liang, Yuanqi Zhai, Shiqi Yang, Qing Gu, Fang Wei, Hong Zhu, Fenghua Wang, Xueting Luo, Xiaodong Sun

## Abstract

Photoreceptor degeneration is a leading cause of visual impairment worldwide. Separation of neurosensory retina from the underlying retinal pigment epithelium is a prominent feature preceding photoreceptor degeneration in a variety of retinal diseases. Although ophthalmic surgeries have been well developed to restore retinal structures, post-op patients usually experience progressive photoreceptor degeneration and irreversible vision loss that is incurable at present. Previous studies point to a critical role of mitochondria-mediated apoptotic pathway in photoreceptor degeneration, but the upstream triggers remain largely unexplored. In this study, we show that after experimental RD induction, photoreceptors activate dynamin-related protein 1 (Drp1)-dependent mitochondrial fission pathway and subsequent apoptotic cascades. Mechanistically, endogenous ROS is necessary for Drp1 activation in vivo and exogenous ROS insult is sufficient to activate Drp1-dependent mitochondrial fission in cultured photoreceptors. Accordingly, inhibition of Drp1 activity effectively preserves mitochondrial integrity and rescues photoreceptors. Collectively, our data delineates a ROS-Drp1-mitochondria axis that promotes photoreceptor degeneration in retinal diseased models.

## Introduction

Photoreceptors are light-sensing neurons responsible for visual signal reception. (Nakanishi, 1995) Due to sustained photo-transduction and oxidative stress,(Ahmed et al., 1993) photoreceptors are among the most metabolically active and energy-demanding tissues in the central nervous system.(Okawa et al., 2008) However, the outer retina, where photoreceptors reside in, mainly depends on retinal pigment epithelium layer and choroid vascular bed underneath for nutrition and oxygen to survive.(Linsenmeier and Padnick-Silver, 2000)’(Luo et al., 2013). Such unique organization makes photoreceptors exceptionally vulnerable to metabolic perturbations under stressed conditions including rhegmatogenous retinal detachment (RD), tractional diabetic retinopathy and age-related macular degeneration.(Arroyo et al., 2005)’(Lim et al., 2012) Clinically palliative interventions remain the mainstay to halt disease progression, but in the long run patients suffer from progressive vision loss due to irreversible photoreceptor death.(Wubben et al., 2016) Therefore, development of photoreceptor-targeted neuroprotective strategy is pivotal to preserve vision.

Photoreceptors degenerate mainly through apoptotic pathway. However, inhibition of death effectors does not effectively protect photoreceptors because of activation of necrotic pathway.(Murakami et al., 2013) Therefore, elucidation of mechanisms leading to photoreceptor degeneration, in particular identification of the initial ‘triggers’ upstream of death effectors, is critical to protect photoreceptors. Nonetheless, the upstream signals of photoreceptor degeneration remain largely unexplored.

Mitochondria are energy-producing organelles that play a central role in cell death decision.(Vakifahmetoglu-Norberg et al., 2017) Accumulating evidence suggests that mitochondrial dysfunction precedes neuronal death in neurodegenerative diseases.(Ishihara et al., 2009) Importantly, decline in mitochondrial metabolism is associated with age-related photoreceptor degeneration,(Kam et al., 2015) indicating the importance of mitochondria for photoreceptor survival. To further clarify the role of mitochondria in photoreceptor degeneration, we employed diseased models both in vivo and in vitro wherein photoreceptors were exposed to stress. We found that mitochondrial fission within photoreceptors represented an early event before activation of death effectors. At the molecular level, we identified Drp1 as the critical mediator of mitochondrial fission in stressed photoreceptors. ROS was both required and sufficient to induce Drp1-dependent mitochondrial fission. Moreover, inhibition of Drp1 substantially preserved mitochondrial integrity and protected photoreceptors both structurally and functionally. Collectively, our findings have uncovered the critical role of mitochondria in photoreceptor degeneration and identified Drp1 as a promising therapeutic target for photoreceptor protection.

## Materials and Methods

### Animals and surgery

The biomedical research was approved by the Shanghai General Hospital review board. All procedures were adhered to the Statement of the Association for Research in Vision and Ophthalmology for using animals in biomedical research. SD rats were housed at the Laboratory Animal Center of Shanghai General Hospital. Male rats (8-10 weeks; 180-250g) were used to conduct retinal detachment as previous description.(Liu et al., 2009) Briefly, the rats were anesthetized by intraperitoneal injection with 1% sodium pentobarbital (Sigma-Aldrich, US). Before surgery, pupils were dilated with 0.5% tropicamide and 0.5% phenylephrine hydrochloride eye drops (Santen Pharmaceutical, Japan). A needle head was used to create a sclerotomy 2mm posterior to the limbus. Then a 30-gauge needle was introduced through the sclerotomy into the subretinal space. Sodium hyaluronate (10 mg/mL, LG Life Sciences, Korea) was injected to make two-thirds of the neurosensory retina detach from the underlying RPE. The RD was verified by surgical microscope and only the right eye was generated to RD, with left eye serving as control. All animal experiments included three to four rats per group and repeated three times.

### Subretinal injections of Mdivi-1 or N-Acetylcysteine

Mdivi-1 (Sigma-Aldrich #338967-87-6) was dissolved in 20% dimethyl sulfoxide (DMSO) and then further diluted with 0.9% physiological saline to a working concentration of 2.4mg/ml. N-Acetylcysteine (Sigma #A7250) was dissolved in 0.9% physiological saline into the working concentration of 200 µM or 100 µM. 5 µL of chemicals or vehicle was injected into subretinal space at the time of RD induction.

### Cell Culture

The 661W cell line was maintained in Dulbecco’s modified Eagle’s medium containing 10% fetal bovine serum, 300 mg/L glutamine, 32 mg/L putrescine, 40 mL/L of b-mercaptoethanol, and 40 mg/L of both hydrocortisone 21-hemisuccinate and progesterone. The media also contained penicillin (90 units/mL) and streptomycin (0.09 mg/mL). Cells were maintained under regular cell culture chamber. 500 µM H_2_O_2_ was used to mimic the oxidative stress induced model for 24 hours.50 µM Mdivi-1 was pretreated to 661 cells for 1 hour before 500 µM H_2_O_2_ treatment.

### Western blot analysis

The retina or cultured cells were homogenized with lysis buffer containing 50 mM Tris 7.4, 150 mM NaCl, 1% Triton X-100, 1% sodium deoxycholate, 0.1% SDS and inhibitors of protease (Roche #11679498001) and phosphatase (Sigma #P0044). Samples were resolved by SDS-PAGE and transferred to PVDF membranes. Primary antibodies used for probing were listed below: phospho-Drp1(Ser616) (Cell Signaling Technology, Product #3455), Drp1 (Cell Signaling Technology, Product #14647), Cleaved Caspase-3 (Proteintech, Product #25546-1-AP), Bax (Abcam, Product #ab32503), VDAC (Cell Signaling Technology, Product #4866), GAPDH (Proteintech, Product #60004) and β-actin (Cell Signaling Technology, Product #4970). After HRP-conjugated secondary antibodies and chemiluminescent substrates were applied, the signals were detected by Amersham Imager 600 (GE, USA). Densitometry was measured and analyzed with Image J 1.48.

### Mitochondrial Fission confocal microscopy

The rats were perfused transcardially with 4% paraformaldehyde. The eyeballs were harvested, cryo-sectioned and prepared for immunohistochemistry. 661W cells were fixed with 4% paraformaldehyde before staining. Leica TCS SP8 confocal microscopy (Leica Microsystems, Weltzlar, Germany) was used to detect V-β antibodies (Abcam #ab14730) labeled mitochondria with an excitation wavelength of 543nm and an emission wavelength from 575nm to 700nm. The image size was set to 1024 x 1024 pixels in a scan speed of 100 Hz with a 63X oil immersion lens. The way of mitochondrial morphology was analyzed from Rehman et al (Rehman et al., 2012).Ten to fifteen cells were captured randomly and the images were analyzed using Image J (NIH,Bethesda, MD). Greater than half of mitochondrial displaying the long tubular shape is “tubular”, less than half of mitochondrial displaying the tubular shape is intermediate and majority of mitochondrial displaying short is fragmented shape.(Wang et al., 2015) Fragmentation is defined as number of the fragmented mitochondrial in total cells.100 cells were statistically analyzed by a blind observer.(Ballweg et al., 2014)

### Histopathological retinal ONL Damage Assessment

Eyes were enucleated 7 days after retinal detachment. Each group consisted of 3 eyes each time and repeated 3 times. Sections (5 µm) were in the vertical meridian and inferior portion of the eye wall and stained with hematoxylin and eosin. Sections in the angle were excluded in our research, because the retinal thickness varies with the distance from the optic nerve, the INL thickness was used to as the internal control at the same distance from the optic nerve(Dong et al., 2012). 10 points in each section were measured by skilled observer, five sections were randomly selected in each eye. The thickness ratio of the ONL to INL was calculated to compare the ONL damage in each group(Besirli et al., 2010)(Trichonas et al., 2010).

### Transmission electron microscopy

TEM analysis was carried out as described previously(Huang et al., 2014). Briefly, the retina was fixed in a solution containing 5% formaldehyde, 2% glutaraldehyde in 0.1 M PBS, pH 7.4. The fixed and processed samples were subjected to imaging with TEM (Zeiss 190). Mitochondrial of photoreceptor were gained randomly in four field for each sample and the images were digitized and the mitochondrial length was measured by Image J 1.48 for analysis.

### Mitochondrial isolation

Mitochondrial isolation from retina was performed using a mitochondrial isolation kit (Beyotime #C3606) according to the instructions. Briefly, dissected retina homogenized in cold mitochondrial lysis buffer. The homogenate was then centrifuged at 600g for 10 min at 4 °C to spin down the nuclei and unbroken cells. The supernatant was collected and centrifuged again at 12,000g for 30 min at 4 °C to pallet the mitochondria.

### Apoptosis assay

DNA fragmentation was detected using terminal deoxynucleotidyl transferase dUTP nick end labeling (TUNEL; Roche #11684795910) 24 hours after H_2_O_2_ treatment. TUNEL-positive staining was accounted in 4 random fields for each group and repeated 3 times.

### Drp1 silencing

661W cells were transfected with siRNA/lipofectamine mixture according to the manufacturer’s instructions (GenePharma, Shanghai, China). We selected four sequences of siRNA from the pubmed and one sequences of siRNAs proved to be effective in this study: siDrp1 sense: 5’-AGGAGAAGAAAAUGGUAAAUUUCTT-3’, siDrp1 antisense: 5’-GAAAUUUACCAUUUUCUUCUCCUTT-3’, negative control sense: 5’-UUCUCCGAACGUGUCACGUTT-3’, negative control antisense: 5’-ACGUGACACGUUCGGAGAATT-3’. The final concentration of siRNA was 20 pM per well. The media was changed 24h after transfection. Cells were harvested 48 h after transfection. Efficiency of knocking down endogenous Mouse Drp1 was verified by Western blot, the transfection efficiency is calculated by the expression of p-Drp1 Ser^616^ between siDRP1 and control group relative to β-actin.

### Mitochondrial membrane potential assay

Dual-emission potential-sensitive probe JC-1 (Molecular Probes #T3168) was used to measure the mitochondrial membrane potential, 661w cells were washing with cold PBS after treatment and then incubated with 10 µL of JC-1 for 20 minutes at 37^°^C according to manufacturer’s instructions. After incubation, cells were washed twice by cold PBS, we measured the red fluorescence with excitation at 525 nm and emission at 590 nm, the green fluorescence with excitation at 490 nm and emission at 530 nm in a random order and analyzed with Image J 1.48. The ratio of red to green fluorescence was used to reflect mitochondrial membrane potential.

### Electroretinography

The retinal function was assessed by electroretinograms (ERG) before and 7 days after retinal detachment. The rats were pretreated overnight dark adaptation, the rats were anesthetized and pupil were dilated as previously introduced. The body temperature was set to 37°C with a heating pad during the procedure, a gold wire electrode was on the corneal, reference electrode was at the head and a ground electrode in the tail. All the procedure was in dim red light. The response to a light flash (3.0 candela seconds/m^2^) from a photic stimulation was amplified, the preamplifier bandwidth was set at 0.2 to 300 Hz. The a-wave amplitude was from baseline to the maximum a-wave. The amplitude of b-wave was measured from the maximum a-wave to the maximum b-wave peak. The ratio of a or b wave to baseline was used to evaluate the retinal function.

### Statistics Analysis

Data were shown as mean ± SD, Difference in 2 groups were analyzed by Unpaired Student’s t-Test, the mitochondrial length, ERG data in three or more groups were analyzed by one-way ANOVA with Bonferroni coefficient, mitochondrial morphology were analyzed by chi-square test by computer software (SPSS 21.0 for windows; SPSS Inc., Chicago, IL). A p value <0.05 was considered as significant.

## Results

### Mitochondrial fission is evident in the photoreceptors after experimental RD

Mitochondria, structurally dynamic organelles, undergoes constant fusion and fission to regulate their homeostasis (Westermann, 2010). Increased mitochondrial fission is associated with functional deficiency under diseased conditions (Archer, 2013). To directly visualize mitochondria morphology in vivo, we employed transmission electron microscopy (TEM) to analyze the retinal tissues of rats on the third day after experimental RD. As shown in Fig. 1A & 1B, the mitochondria within photoreceptors appeared to be fragmented, punctate and scattered as compared to the typically long and interconnected structures in untreated control. We also labeled the mitochondria with antibodies against COX-V β subunit (V-β) for analysis of mitochondrial morphology with confocal microscopy. In normal photoreceptors, rod-like structures in various lengths were readily detectable. However, punctate and dispersed mitochondrial signals dominated the diseased photoreceptors after RD (Fig. 1C & 1D). We artificially divided the mitochondria into three species (i.e. long, intermediate and tubular) based on their morphology and further assessed the distribution of mitochondrial structures (Picard et al., 2013). Consistent with previous TEM analysis, the amount and proportion of short mitochondrial species increased substantially with concurrent decrease of intermediate and long species moderately in photoreceptors after RD (Fig. 1C & 1D).

**Fig. 1.**
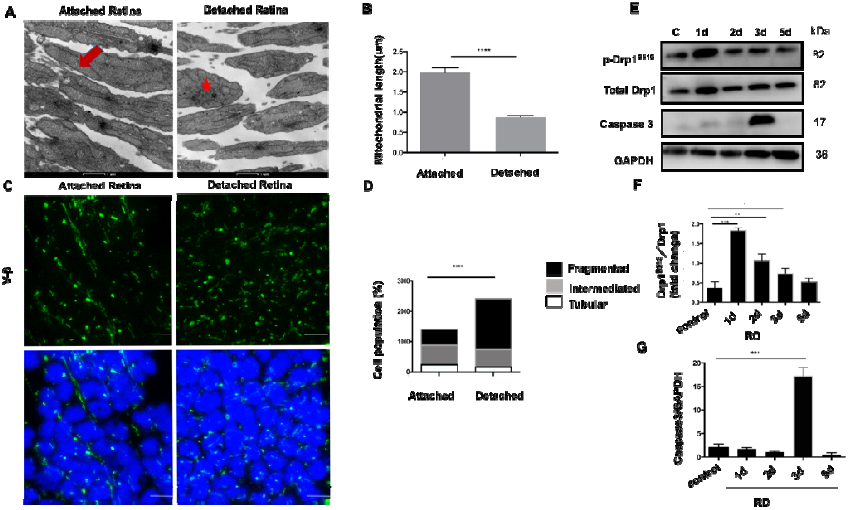
Mitochondrial dynamic changes in the detached retina. (A) Representative TEM images showed mitochondrial network of photoreceptors in attached retina and detached retina 3days after RD. Arrow, star mark tubular and fragmented mitochondrial species, respectively; Scale bar: 1µm. **(B)** Quantification of mitochondrial length in photoreceptors, mitochondrial length become shorter after RD, \*\*\*\**P*<0.0001, independent t test, 3 independent experiments. **(C)** Mitochondrial network of the photoreceptors was labeled with V-β antibodies and assessed by confocal microscopy in attached and detached retina 3 days after RD. Greater than half of mitochondrial displaying the long tubular shape is “tubular”, less than half of mitochondrial displaying the tubular shape is intermediate and majority of mitochondrial displaying short is fragmented shape; Scale bars: 5um. **(D)** Quantification of mitochondrial morphology by Image J, (cell number =100), the fragmentation was defined as the percentage of fragmented mitochondrial in total cells, fragmented mitochondrial ratio increased after RD, \*\*\*\**P* <0.0001, Chi-square test, 3 independent experiments. **(E-G)** Western blot analysis of retinal lysates after RD (n=3 for each group), quantification indicates that mitochondrial fission protein Drp1S^616^ increased and peaked 1 day after RD, \*\*\*\**P* <0.0001, caspase-3 increased and peaked 3 days after RD, \*\*\*\**P* <0.0001, ANOVA with Bonferroni coefficient, 3 independent experiments.

### Drp1 activation is critical to mitochondrial fission and subsequent photoreceptor death

Drp1 is the essential mediator of mitochondrial fission and its phosphorylation is associated with activation of the mitochondrial fission pathway (Kashatus et al., 2015). In order to assess whether Drp1 is involved in the process of mitochondrial fission in photoreceptors after RD, we dissected out the retina for analysis by Western blot. As showed in Fig. 1E and Fig.1F, Drp1 appeared to be phosphorylated abruptly after injury. The p-Drp1 signal peaked within one day after RD followed by gradual decline, suggesting a potential role of Drp1 in RD-induced mitochondrial fission in photoreceptors. Thus, We conclude that RD induced Drp1 activation and mitochondrial fission in the photoreceptors.

### Inhibition of Drp1 attenuates mitochondrial fission and photoreceptor degeneration after RD

Apoptosis is the primary pathway through which photoreceptors degenerate (Murakami et al., 2013). Consistently, we detected significant up-regulation of Caspase-3 activity at 3 days after RD (Fig. 1E & 1G). Since activation of Drp1 preceded expression of Caspase-3 (Fig. 1F & 1G), we imply that Drp1-mediated mitochondrial fission may serve as upstream signals regulating the apoptotic pathway in photoreceptors after RD. To further verify this hypothesis, we subretinally administered Mdivi-1, a highly selective inhibitor for Drp1 (Tanaka and Youle, 2008), at the time of RD induction. As expected, treatment with Mdivi-1 attenuated RD-induced mitochondrial fission and effectively preserved mitochondrial integrity in the photoreceptors after RD as determined by TEM (Fig. 2A & 2B). Next, we analyzed the activity of apoptotic factors in the retinal tissues. Notably, we analyzed the activity of apoptotic factors in the retinal tissues. Notably, the expression of cleaved Caspase-3 was substantially suppressed by Mdivi-1 after RD (Fig. 2C & 2D). The Bcl-2 family is a critical regulator of mitochondrial permeability and intrinsic apoptotic pathway (Chipuk et al., 2010). Previous report has demonstrated a critical role of Bax, a member of the Bcl-2 family, in photoreceptor degeneration after RD (Yang et al., 2004). To test whether Bax is involved in Mdivi-1 mediated suppression of Caspase-3 activity, we further analyzed the expression level of Bax in the retinal lysates after RD. As shown in Fig. 2C & 2D, Bax expression was upregulated significantly after RD, while Mdivi-1 treatment substantially attenuated Bax activity. Both Drp1 and Bax are cytoplasmic proteins that translocate onto mitochondria membrane upon activation (Frank et al., 2001), (Große et al., 2016). To this end, we extracted the mitochondrial fraction from the retinal lysate and found that both activated Drp1 and Bax protein in mitochondria were substantially attenuated (Fig. 2E & 2F).

**Fig. 2.**
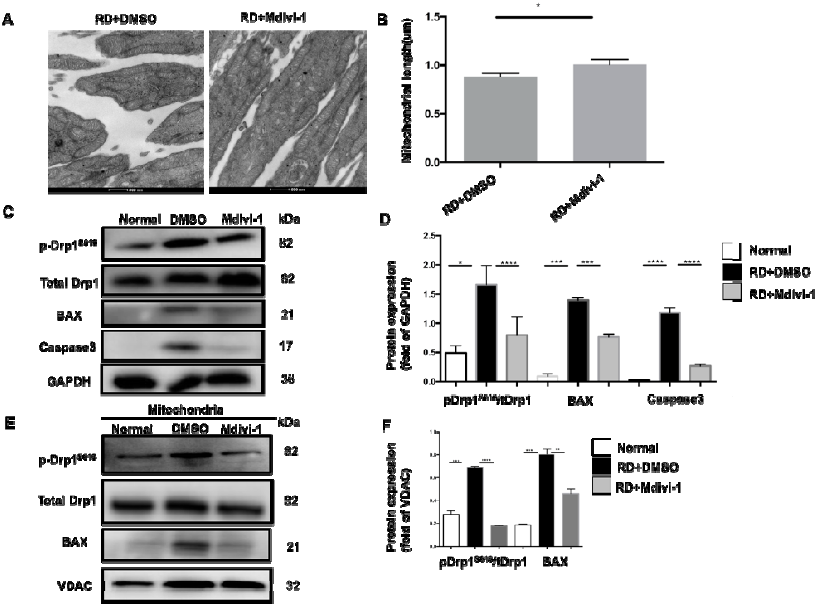
Inhibition of Drp1 protects the photoreceptors by suppressing intrinsic apoptosis way. **(A-B)** Representative TEM images of the mitochondria of photoreceptors in RD and RD treated group at 3 days after RD, the mitochondrial length in treated group is longer than DMSO group; Scale bar: 500nm; \**P*=0.026, independent t test, 3 independent experiments. **(C-D)** Western blot analysis of the retinal lysates with Mdivi-1 treatment 3 days after RD (n=3), GAPDH was used as internal control for analysis, Drp1S^616^/tDrp1, caspas3 and Bax decreased with Mdivi-1 treatment, \*\*\*\**P*<0.0001, ANOVA with Bonferroni coefficient, 3 independent experiments; **(E-F)** Western blot analysis of the mitochondrial fraction with Mdivi-1 treatment 3 days after RD (n=3). VDAC was used as a marker for mitochondria and as internal control. Quantification indicates that p-Drp1, BAX decreased compared with DMSO group; \*\*\*\**P*<0.0001, \*\*\**p*=0.0046, ANOVA with Bonferroni coefficient, 3 independent experiments.

Based on the inhibitory effect of Mdivi-1 on the mitochondrial fission and apoptotic pathways, we further assessed the therapeutic potential of Mdivi-1 in experimental RD model. After RD, progressive degeneration of photoreceptors results in a gradual thinning of the outer nuclear layer (ONL). Therefore, the thickness of ONL serves as an indicator of photoreceptor survival. As showed in Fig. 3A and 3B, RD induced substantial loss of photoreceptors while Mdivi-1 treatment effectively preserved the ONL structure in the retina. Consistently, the retinal function was significantly rescued with Mdivi-1 treatment as examined by scotopic electroretinogram (Fig. 3C, 3D).

**Fig. 3.**
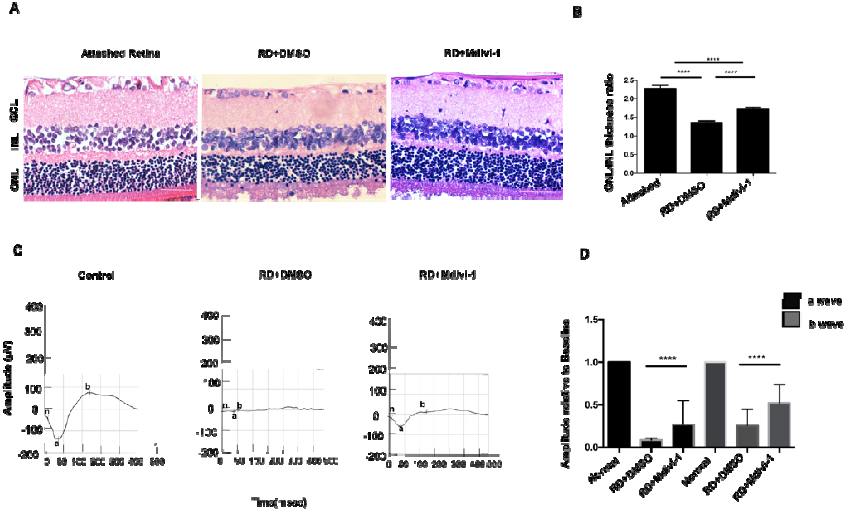
Inhibition of Drp1 attenuated retina function after retinal detachment. **(A-B)** Representative H&E stained retinal sections showed outer segment of the retina 7dasys after RD. The thickness of ONL normalized to that of INL was used as indication of photoreceptor survival. Mdivi-1 treatment group preserved the ONL/INL thickness ratio 7 days after RD (n=3), 3 independent experiments, \*\**P*=0.0061 using ANOVA with Bonferroni coefficient. Scale bars: 50um; **(C-D)** Mdivi-1 treated group increases the functional recovery of scotopic electroretinagram 7 days after RD, **C** representatives scotopic electroretinagram waveforms from baseline, DMSO and Mdivi-1 treated group (n=3). The a or b wave relative to baseline increased after Mdivi-1 treatment, \*\*\*\**P*<0.001, using ANOVA with Bonferroni coefficient, 3 independent experiments.

Taken together, we conclude that Drp1 inhibition by Mdivi-1 has a neuroprotective impact on the retina in experimental RD model by suppressing mitochondrial fission and the apoptotic pathway.

### Drp1-mediated mitochondrial fission is induced by oxidative stress

Oxidative stress has been well documented as a ‘danger signal’ to photoreceptors. Accordingly, alleviation of oxidative stress protects photoreceptors after RD (Roh et al., 2011). Previous studies indicated potential correlations between oxidative stress and mitochondrial fission (Yu et al., 2014). However, their causal relationship remains to be explored. Therefore, we questioned whether attenuation of oxidative stress would have an effect on mitochondrial dynamics in photoreceptors after RD. To this end, we introduced N-Acetylcysteine (NAC), a well-defined scavenger of reactive oxygen species (ROS), subretinally at the time of RD induction and examined the expression level of activated Drp1. As showed in Fig. 4A-4B, NAC effectively suppressed the expression of p-Drp1, indicating a positive role of oxidative stress in promoting mitochondrial fission.

**Fig. 4.**
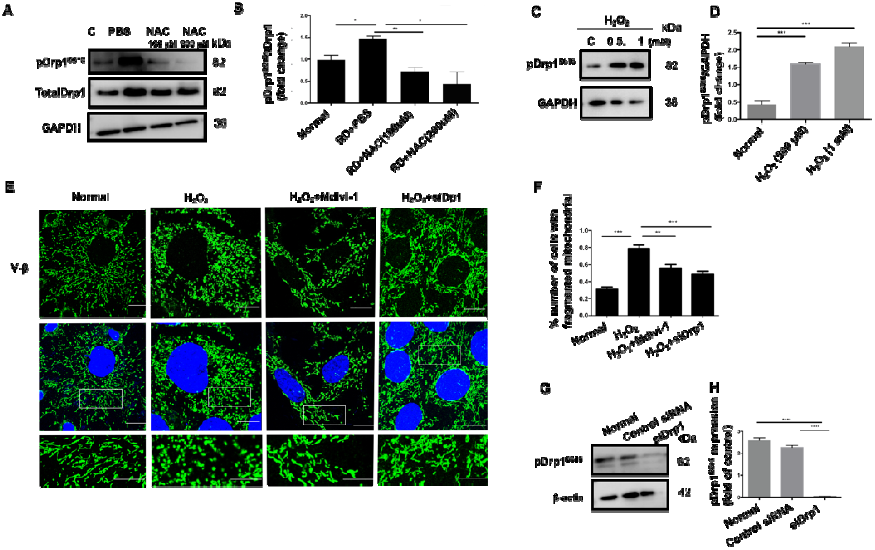
Oxidative stress induced mitochondrial fission. **(A-B)** Western blot analysis of pDrpS^616^/tDrp1 in retinal lysates 3 days after RD and NAC treatment(n=3). Quantification results indicate that NAC treated group decreased the pDrpS^616^/tDrp1 expression, GAPDH was used as internal control, \**P*=0.0212, \*\*\**P*=0.0031, \**P* =0.0213, using ANOVA with Bonferroni coefficient, 3 independent experiments. **(C-D)** Representative western blot of pDrpS^616^ in different H_2_O_2_ concentrations insulted 661W cells for 24 hours; Quantification results indicate that pDrpS^616^ increased after H_2_O_2_ treatment, GAPDH was used as internal control; \*\*\**P*=0.002, \*\*\**P*=0.002, respectively, 3 independent experiments. **(E-F)** 500 µM H_2_O_2_ was induced in 661W cells for 24 hours, morphology of mitochondria was imaged by confocal microscopy and quantitated with Image J (n=100). H_2_O_2_ increased mitochondrial fragmentation, \*\*\*\**P*<0.0001 for H_2_O_2_ VS normal group, however Mdivi-1 or siDrp1 reduced the effect, \*\**P*=0.0052 for H_2_O_2_ VS Mdivi-1 treated group, \*\*\**P*<0.0001 H_2_O_2_ VS for siDrp1, Chi-square test. Scale bar: 10um. 3 independent experiments. **(G-H)** Representative western blot of siDrp1 after transfection; Quantification of p-Drp1Ser^616^ normalized to β-actin in three groups (n=3), siDrp1 group decreased the p-Drp1Ser^616^ expression; \*\*\*\**P*<0.0001, ANOVA with Bonferroni coefficient, the transfection efficiency is 91.5%, 3 independent experiments.

To further elucidate the causal relationship between oxidative stress and mitochondrial fission, we employed an in vitro model of cultured murine photoreceptor-derived 661W cell line challenged by H_2_O_2_. Consistent with what we found in vivo, H_2_O_2_ insult activated Drp1 as showed in Fig. 4C & 4D. Immunostaining of 661W cells with mitochondria-specific V-β antibodies revealed predominance of fragmented mitochondrial signals after H_2_O_2_ insult (Fig. 4E & 4F), which is in agreement with our vivo findings (Fig. 4A). Consistently, Drp1 knockdown by siRNA (Fig. 4G & 4H) also preserved mitochondrial integrity (Fig. 4E & 4F), confirming the critical role of Drp1.

### Drp1 inhibition suppresses oxidative stress induced degenerative signals

H_2_O_2_ insult upregulated intracellular ROS level of cultured 661W cells and cell degeneration, while, inhibition of Drp1 activity by Mdivi-1 or siRNAs effectively suppressed H_2_O_2_ induced TUNEL activity (Fig. 5A & 5B), Pre-treatment of 661W cells with Mdivi-1 effectively preserved mitochondrial integrity and membrane potential (Fig. 5C-5D), which supports a role of Drp1 in oxidative stress induced mitochondrial dysfunction. Our findings strongly suggest that Drp1 and Drp1-dependent pathways may join in preserving the mitochondrial function, Oxidative stress induced photoreceptor degeneration is at least in part mediated by Drp1-dependent mitochondrial fission.

**Fig. 5.**
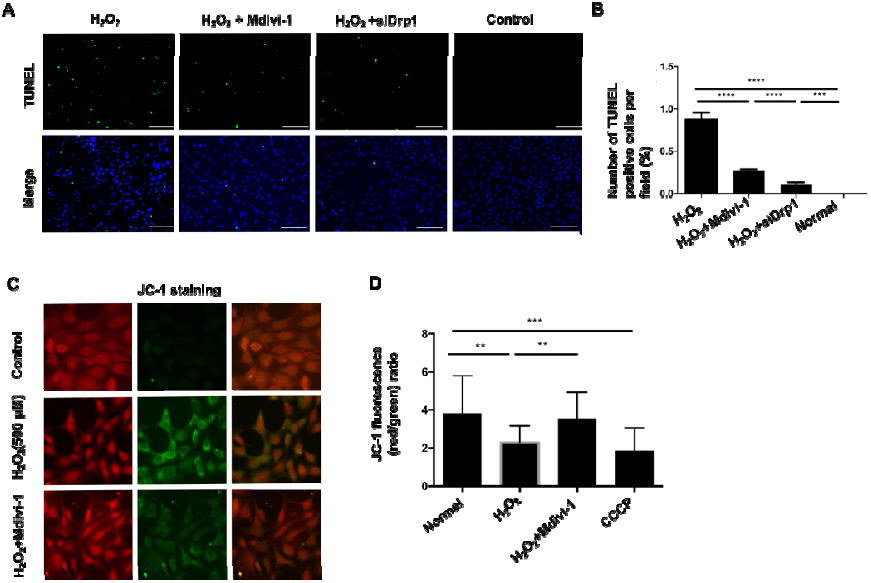
Inhibition of Drp1 suppresses apoptosis and protects 661W cells through protecting mitochondrial integrity after H_2_O_2_ insult. **(A-B)** Representative TUNEL staining after treatment, either Mdivi-1 or siDrp1 was effective in rescuing 661W cells after 500 µM H_2_O_2_ insult for 24 hours. Quantification of TUNEL positive cells in total cells, \*\*\**P*=0.0002 for Mdivi-1 VS H_2_O_2_, \*\*\**P*=0.0008 for siDrp1 VS H_2_O_2_, ANOVA with Bonferroni coefficient, 3 independent experiments. **(C-D)** Representative mitochondrial membrane potential by JC-1 staining in different groups. JC-1 dye expresses red signals with high membrane potential and shifts to green fluorescence when mitochondria was decoupled. Quantification of red and green fluorescence intensity and calculated the red/green ratio after JC-1 staining in the H_2_O_2_ induced model, Mdivi-1 rescued H_2_O_2_-induced loss of mitochondrial membrane potential and increased the red/green ratio. Scale bar: 10um. \*\**P*=0.0011, ANOVA with Bonferroni coefficient. 3 independent experiments.

### Discussion

Photoreceptors play pivotal roles in visual reception by a number of energy-demanding activities including phototransduction, resolution of light and oxygen-induced damage, synthesis and replenishment of disc membranes, etc (Ahmed et al., 1993). Therefore, photoreceptors necessitate sustained supply of oxygen and nutrition from the RPE/choroid complex to support highly active metabolism (Linsenmeier and Padnick-Silver, 2000), (Luo et al., 2013). Accordingly, photoreceptors hold a large reservoir of mitochondria to satisfy their metabolic demand(Barber and Wright, 1969). It has been reported that increased mitochondrial fission is associated with photoreceptor degeneration in aged mice(Kam et al., 2015). Moreover, mitochondrial abnormality correlates with photoreceptor degeneration in acute glucose deprivation model.(Chertov et al., 2011) However, the relationship between mitochondrial abnormality and photoreceptor death remains unknown.

In the case of RD as manifested in multiple retinal diseases, photoreceptors suffer from excessive ROS that is considered to trigger cell death, but the exact mechanisms remain to be elucidated. Accumulating evidence suggests that mitochondria is the major source of intracellular ROS in response to cellular stress(Sies, 2014). Consistently, we observed a predominantly fissured population of mitochondria in the photoreceptors of rats after experimental RD, which represents an early degenerative event (Fig.1E & 1F). Since fissured mitochondria has been associated with energetic failure and cell degeneration(Cherubini and Ginés, 2017), we speculate that mitochondrial fission could be the milestone along the ROS-induced degenerative pathway determining photoreceptor death. At the molecular level, Drp1, the key mediator of mitochondrial fission, was activated after RD. Notably, inhibition of ROS suppressed Drp1 activation, which supports a regulatory role of ROS in Drp1 activity. In vitro, H_2_O_2_ induced Drp1 activation, mitochondrial fission and photoreceptor death, while inhibition of Drp1 preserved mitochondrial integrity and protected photoreceptors. Collectively, our data suggest that Drp1-dependent mitochondrial fission plays key roles in mediating RD-induced photoreceptor degeneration.

It is intriguing to target mitochondria for photoreceptor protection since mitochondria are considered to be the major organelles that senses cellular stress and broadcasts death-promoting signals. Mitochondrial fission may represent a common pathway through which photoreceptors degenerate in other retinal diseased models. In conclusion, our findings suggest that Drp1-dependent mitochondrial fission plays critical roles in photoreceptor degeneration and Drp1 inhibition is effective in preserving photoreceptors. Thus, Drp1 represents a potential therapeutic target for photoreceptor protection.

## Acknowledgments

We thank Dr. Yu Chen (Yueyang Hospital, Shanghai University of Traditional Chinese Medicine) for providing the 661W photoreceptor cell line.

## References

Ahmed, J., Braun, R. D., Dunn, R. and Linsenmeier, R. A. (1993).Oxygen distribution in the macaque retina. Investigative Ophthalmology and Visual Science 34, 516–521.

Archer, S. L. (2013).Mitochondrial Dynamics - mitochondrial fission and fusion in human diseases. New England Journal of Medicine 369, 2236–2251.

Arroyo, J. G., Yang, L., Bula, D. and Chen, D. F. (2005).Photoreceptor apoptosis in human retinal detachment. American journal of ophthalmology 139, 605–10.

Ballweg, K., Mutze, K., Königshoff, M., Eickelberg, O. and Meiners, S. (2014).Cigarette smoke extract affects mitochondrial function in alveolar epithelial cells. American journal of physiology. Lung cellular and molecular physiology 307, L895–907.

Barber, V. C. and Wright, D. E. (1969).The fine structure of the eye and optic tentacle of the mollusc Cardium edule. Journal of ultrastructure research 26, 515–528.

Besirli, C. G., Chinskey, N. D., Zheng, Q. D. and Zacks, D. N. (2010).Inhibition of retinal detachment-induced apoptosis in photoreceptors by a small peptide inhibitor of the Fas receptor. Investigative Ophthalmology and Visual Science 51, 2177–2184.

Bhatt, L., Groeger, G., McDermott, K. and Cotter, T. G. (2010).Rod and cone photoreceptor cells produce ROS in response to stress in a live retinal explant system. Molecular vision 16, 283–293.

Chertov, A. O., Holzhausen, L., Kuok, I. T., Couron, D., Parker, E., Linton, J. D., Sadilek, M., Sweet, I. R. and Hurley, J. B. (2011).Roles of glucose in photoreceptor survival. Journal of Biological Chemistry 286, 34700–34711.

Cherubini, M. and Ginés, S. (2017).Mitochondrial fragmentation in neuronal degeneration: Toward an understanding of HD striatal susceptibility. Biochemical and Biophysical Research Communications 483, 1063–1068.

Chipuk, J. E., Moldoveanu, T., Llambi, F., Parsons, M. J. and Green, D. R. (2010).The BCL-2 Family Reunion. Molecular Cell 37, 299–310.

Dong, K., Zhu, H., Song, Z., Gong, Y., Wang, F., Wang, W., Zheng, Z., Yu, Z., Gu, Q., Xu, X., et al. (2012).Necrostatin-1 protects photoreceptors from cell death and improves functional outcome after experimental retinal detachment. American Journal of Pathology 181, 1634–1641.

Frank, S., Gaume, B., Bergmann-Leitner, E. S., Leitner, W. W., Robert, E. G., Catez, F., Smith, C. L. and Youle, R. J. (2001).The Role of Dynamin-Related Protein 1, a Mediator of Mitochondrial Fission, in Apoptosis. Developmental Cell 1, 515–525.

Große, L., Wurm, C. A., Brüser, C., Neumann, D., Jans, D. C. and Jakobs, S. (2016).Bax assembles into large ring-like structures remodeling the mitochondrial outer membrane in apoptosis. The EMBO journal 35, 402–413.

Huang, Q., Li, J., Xing, J., Li, W., Li, H., Ke, X., Zhang, J., Ren, T., Shang, Y., Yang, H., et al. (2014).CD147 promotes reprogramming of glucose metabolism and cell proliferation in HCC cells by inhibiting the p53-dependent signaling pathway. Journal of Hepatology.

Ishihara, N., Nomura, M., Jofuku, A., Kato, H., Suzuki, S. O., Masuda, K., Otera, H., Nakanishi, Y., Nonaka, I., Goto, Y., et al. (2009).Mitochondrial fission factor Drp1 is essential for embryonic development and synapse formation in mice. Nature Cell Biology 11, 958–966.

Kam, J. H., Jeffery, G., Hoh Kam, J. and Jeffery, G. (2015).To unite or divide: mitochondrial dynamics in the murine outer retina that preceded age related photoreceptor loss. Oncotarget 6, 26690–26701.

Kashatus, J. A., Nascimento, A., Myers, L. J., Sher, A., Byrne, F. L., Hoehn, K. L., Counter, C. M. and Kashatus, D. F. (2015).Erk2 phosphorylation of Drp1 promotes mitochondrial fission and MAPK-driven tumor growth. Molecular Cell 57, 537–552.

Lim, L. S., Mitchell, P., Seddon, J. M., Holz, F. G. and Wong, T. Y. (2012).Age-related macular degeneration. The Lancet 379, 1728–1738.

Linsenmeier, R. A. and Padnick-Silver, L. (2000).Metabolic dependence of photoreceptors on the choroid in the normal and detached retina. Investigative Ophthalmology and Visual Science 41, 3117–3123.

Liu, H., Qian, J., Wang, F., Sun, X., Xu, X., Xu, W. and Zhang, X. (2009).Expression of two endoplasmic reticulum stress markers, GRP78 and GADD153, in rat retinal detachment model and its implication. Eye 24, 137–144.

Luo, L., Uehara, H., Zhang, X., Das, S. K., Olsen, T., Holt, D., Simonis, J. M., Jackman, K., Singh, N., Miya, T. R., et al. (2013).Photoreceptor avascular privilege is shielded by soluble VEGF receptor-1. eLife 2013,.

Murakami, Y., Notomi, S., Hisatomi, T., Nakazawa, T., Ishibashi, T., Miller, J. W. and Vavvas, D. G. (2013).Photoreceptor cell death and rescue in retinal detachment and degenerations. Progress in Retinal and Eye Research 37, 114–140.

Nakanishi, S. (1995).2nd-Order Neurons and Receptor Mechanisms in Visual-Information and Olfactory-Information Processing. Trends in Neurosciences 18, 359–364.

Okawa, H., Sampath, A. P., Laughlin, S. B. and Fain, G. L. (2008).ATP Consumption by Mammalian Rod Photoreceptors in Darkness and in Light. Current Biology 18, 1917–1921.

Picard, M., Shirihai, O. S., Gentil, B. J. and Burelle, Y. (2013).Mitochondrial morphology transitions and functions: implications for retrograde signaling AJP: Regulatory, Integrative and Comparative Physiology 304, R393–R406.

Rehman, J., Zhang, H. J., Toth, P. T., Zhang, Y., Marsboom, G., Hong, Z., Salgia, R., Husain, A. N., Wietholt, C. and Archer, S. L. (2012).Inhibition of mitochondrial fission prevents cell cycle progression in lung cancer. The FASEB Journal 1–12.

Roh, M. I., Murakami, Y., Thanos, A., Vavvas, D. G. and Miller, J. W. (2011).Edaravone, an ROS scavenger, ameliorates photoreceptor cell death after experimental retinal detachment. Investigative Ophthalmology and Visual Science 52, 3825–3831

Sies, H. (2014).Role of metabolic H2O2 generation: Redox signaling and oxidative stress. Journal of Biological Chemistry 289, 8735–8741.

Tanaka, A. and Youle, R. J. (2008).A Chemical Inhibitor of DRP1 Uncouples Mitochondrial Fission and Apoptosis. Molecular Cell 29, 409–410.

Trichonas, G., Murakami, Y., Thanos, A., Morizane, Y., Kayama, M., Debouck, C. M., Hisatomi, T., Miller, J. W. and Vavvas, D. G. (2010).Receptor interacting protein kinases mediate retinal detachment-induced photoreceptor necrosis and compensate for inhibition of apoptosis. Proceedings of the National Academy of Sciences of the United States of America 107, 21695–21700.

Vakifahmetoglu-Norberg, H., Ouchida, A. T. and Norberg, E. (2017).The role of mitochondria in metabolism and cell death. Biochemical and Biophysical Research Communications 482, 426–431.

Wang, L., Yu, T., Lee, H., O’Brien, D. K., Sesaki, H. and Yoon, Y. (2015).Decreasing mitochondrial fission diminishes vascular smooth muscle cell migration and ameliorates intimal hyperplasia. Cardiovascular Research 106, 272–283.

Westermann, B. (2010).Mitochondrial fusion and fission in cell life and death. Nature reviews. Molecular cell biology 11, 872–84.

Wubben, T. J., Besirli, C. G. and Zacks, D. N. (2016).Pharmacotherapies for Retinal Detachment. Ophthalmology 123, 1553–1562.

Yang, L., Bula, D., Arroyo, J. G. and Chen, D. F. (2004).Preventing Retinal Detachment-Associated Photoreceptor Cell Loss in Bax-Deficient Mice. Investigative Ophthalmology and Visual Science 45, 648–654.

Yu, T., Wang, L., Lee, H., O’Brien, D. K., Bronk, S. F., Gores, G. J. and Yoon, Y. (2014).Decreasing mitochondrial fission prevents cholestatic liver injury. Journal of Biological Chemistry 289, 34074–34088.

